# Bed Nucleus of the Stria Terminalis (BNST) neurons containing the serotonin 5HT_2c_ receptor modulate operant alcohol self-administration behavior in mice

**DOI:** 10.1101/2023.09.26.559653

**Authors:** Meghan E. Flanigan, Carol Gianessi, Megan Castle, Winifred Dorlean, Tori Sides, Thomas L. Kash

## Abstract

The serotonin 5HT_2c_ receptor has been widely implicated in the pathophysiology of alcohol use disorder (AUD), particularly alcohol seeking and the affective consequences of chronic alcohol consumption. However, little is known about the brain sites in which 5HT_2c_ exerts its effects on specific alcohol-related behaviors, especially in females. Here, we investigated the effects of site-specific manipulation of the 5HT_2c_ receptor system in the BNST on operant alcohol self-administration behaviors in adult mice of both sexes, including the acquisition and maintenance of fixed-ratio responding, motivation for alcohol (progressive ratio), and quinine-adulterated responding for alcohol on a fixed-ratio schedule (punished alcohol seeking). Knockdown of 5HT_2c_ in the BNST did not affect the acquisition or maintenance of operant alcohol self-administration, nor did it affect progressive ratio responding for alcohol. This manipulation had only a subtle effect on responding for quinine alcohol selectively in females. On the other hand, chemogenetic inhibition of BNST 5HT_2c_-containing neurons (BNST^5HT2c^) increased operant alcohol self-administration behavior in both sexes on day 2, but not day 9, of testing. It also increased operant responding for 1000 μM quinine-adulterated alcohol selectively in males. Importantly, chemogenetic inhibition of BNST^5HT2c^ did not alter operant sucrose responding or motivation for sucrose in either sex. We then performed cell-type specific anterograde tracing, which revealed that BNST^5HT2c^ project to similar regions in males and females, many of which have been previously implicated in AUD. We next used chemogenetics and quantification of the immediate early gene cFos to characterize the functional influence of BNST^5HT2c^ inhibition on vlPAG activity. We show that chemogenetic inhibition of BNST^5HT2c^ reduces vlPAG cFos in both sexes, but that this reduction is more robust in males. Together these findings suggest that BNST^5HT2c^ neurons, and to a small extent the BNST 5HT_2c_ receptor, serve to promote aversive responses to alcohol consumption, potentially through sex-dependent disinhibition of vlPAG neurons.

## Introduction

Alcohol Use Disorder (AUD) is characterized by increased motivation for alcohol and impaired control over alcohol use despite negative consequences. In 2019, 5.6 percent of adults in the U.S. had AUD (NIAAA, 2020), and the prevalence is increasing [1], particularly in females [2]. Thus, understanding the neurobiological mechanisms mediating alcohol seeking, consumption, reward, and aversion is more urgent now than ever.

The brain’s serotonin (5HT) system plays a key role in stress, affect, learning, and reward, making it a target for mediating multiple aspects of AUD [3, 4]. 5HT is involved in both alcohol seeking and its corresponding negative emotional consequences, but its role is complex. 5HT circuitry is extensive throughout the brain, posing a challenge with isolating the contributions of specific brain regions to alcohol-related behaviors (Marcinkiewcz et al., 2016). Adding to the complexity of 5HT neuronal connectivity, the brain expresses a wide variety of 5HT receptors, each with distinct anatomical expression, subcellular localization, and functional downstream effectors.

The bed nucleus of the stria terminalis (BNST) is a functionally and anatomically heterogeneous part of the extended amygdala that has been previously implicated in alcohol-related behaviors, particularly alcohol consumption and alcohol-induced emotional dysregulation. The BNST expresses a wide variety of 5HT receptors, including 5HT_2c_, 2a, 1a, 1b, and 7 [4–6], each of which can play distinct roles in behavior. For example, 5HT_1a_ activation in the BNST reduces anxiety-like behavior [7], while 5HT_2c_ activation promotes it [8]. Though few studies have explored the role of the BNST 5HT system in alcohol-related behaviors, the BNST 5HT_2c_ receptor has been identified as a potential regulator of both alcohol consumption and alcohol’s deleterious effects on socio-affective functioning. Recent studies reported that 5HT_2c_ in the BNST may play a role in alcohol-induced social impairments [9, 10]. With regard to alcohol consumption, genetic deletion of 5HT_2c_ in the BNST selectively increased home-cage alcohol consumption in females, but not males [9]. Consistent with this, chemogenetic activation of 5HT_2c_-containing neurons (BNST^5HT2c^) reduced home-cage binge alcohol consumption in females. Interestingly, there was no effect of chemogenetic inhibition of BNST^5HT2c^ on home cage binge alcohol consumption in either sex. Together, these results suggested that both the 5HT_2c_ receptor and the activation of BNST^5HT2c^ neurons could be important for suppressing alcohol consumption in females.

To further explore the behavioral role of the BNST 5HT_2c_ receptor and BNST^5HT2c^ neurons on volitional operant alcohol consumption-related behaviors, here we employed a model of alcohol self-administration. Specifically, we tested whether genetic knockdown of 5HT_2c_ in BNST or chemogenetic inhibition of BNST^5HT2c^ influenced operant fixed-ratio alcohol seeking, motivation for alcohol (progressive ratio), and quinine-adulterated alcohol seeking (punished alcohol seeking) in adult male and female mice.

## Materials and Methods

### Animals

Adult (aged 2-5 months) male and female 5HT2c-cre mice [11] and 5HT2c^lox/lox^ mice [12] were used for this study. All groups were age-matched for all experiments. Mice were bred in-house and group-housed with littermates (experimental groups were counter-balanced for litter effects). Mice were kept on a reverse light-dark schedule with lights off at 7am and on at 7pm. Mice had ad-libitum access to food (Prolab Isopro RMH 3000, LabDiet) and water until three days prior to behavioral experiments. At this point, mice were maintained at 85- 90% of their body weight through mild food restriction but were given free access to water throughout testing. Mice were fed immediately prior to operant testing each day. All experiments were approved by the [Author University] Institutional Animal Care and Use Committee (IACUC) and are in accordance with the NIH guidelines for the care and use of laboratory animals.

### Surgery

Mice were anesthetized with isoflurane (1-3%) in oxygen (1-2 L/min) and positioned in a stereotaxic frame using ear cup bars (Kopf Instruments). The scalp was sterilized with 70% ethanol and betadine and a vertical incision was made before using a drill to burr small holes in the skull directly above the injection targets. Using a 1 µl Neuros Hamilton Syringe (Hamilton, Inc.), viruses were then microinjected at a 0° angle into the BNST (mm relative to bregma: AP: + 0.7, ML: ± 0.9, DV: -4.55) at a volume of 150 nl of virus per injection site.

For chemogenetic inhibition, 5HT2c-cre mice were injected bilaterally with either AAV8-hSyn-DIO-mCherry, or AAV8-hSyn-DIO-hm4Di-mCherry (Addgene). For 5HT_2c_ knockdown, 5HT2c^lox/lox^ mice were injected bilaterally with either AAV8-hSyn-GFP or AAV8-hSyn-Cre-GFP (UNC Vector Core). For anterograde tracing of 5HT2c- containing BNST neuron synaptic targets, 5HT2c-cre mice were injected in the BNST bilaterally with AAV8-Ef1a- DIO-synaptophysin-mCherry (MIT Vector Core). Mice were given acute ketoprofen subcutaneously (0.1 mg/kg) on the day of surgery and daily for the two days post-op. Mice were allowed to recover in their home cages with ad-libitum food access for at least 4 weeks before the start of behavioral experiments.

### Apparatus

Behavior was assessed in standard operant conditioning chambers placed inside of sound-attenuating boxes (Med Associates). Each chamber was equipped with two levers (active or inactive) located on the right wall, with a reward magazine positioned on the left wall. A fan provided ventilation and background noise throughout the behavioral sessions. Liquid reinforcers (9% ethanol v/v + 2% sucrose w/v or 9% ethanol v/v + 250-1000 µM quinine) were delivered by syringe pump into the magazine cup at a volume of 14 µl per reward. When a reward was delivered, a cue light illuminated above the active lever.

### Fixed-Ratio Alcohol Seeking

The day before training began, mice were exposed to the ethanol + sucrose solution (9% EtOH + 2% sucrose) in their home cage overnight. Active Llver presses were initially reinforced using a fixed ratio 1 (FR1) schedule, where each active lever press resulted in the delivery of a single alcohol reward and the illumination of a cue light. Mice also had the option to press an inactive lever, which did not result in reward or cue delivery. Once individual mice reached a criterion of 10 lever presses on the FR1 schedule, they advanced to an FR2 schedule (2 active lever presses/reward). Mice that did not reach criteria on FR1 after 5 days of testing were excluded from the study. After three days of testing on an FR2 schedule, experimental mice advanced to an FR4 schedule (4 active lever presses/reward) and were tested for 8 days. Each operant session lasted 45 minutes. Measures recorded during testing included: active lever presses, inactive lever presses, rewards, magazine entries, and volume of ethanol left in the magazine at the end of the session (to calculate g/kg consumed).

*Chemogenetic inhibition during fixed-ratio alcohol seeking:* On the second day of FR4 testing, all mice were given 3 mg/kg CNO in saline intraperitoneally (i.p.) 30 minutes prior to their session.

### Progressive Ratio Testing

Following FR4 testing, mice were tested in an exponential progressive ratio test, (single 45 min session) where the number of lever presses required to earn a single ethanol reward exponentially increased over the session according to the formula (rounded to the nearest integer) = [5*e*^(R*0.2)^]-5, where R is equal to the number of rewards already earned plus 1 (i.e., next reinforcer). Thus, the number of responses required to earn a single reward followed the order 1, 2, 4, 6, 9, 12, 15, 20, 25, 32, 40, 50, 62…and so on. The total number of rewards each animal was willing to work for under this schedule was recorded as the “break point.

*Chemogenetic inhibition during progressive ratio testing:* All mice were given 3 mg/kg CNO in saline i.p. 30 minutes before the progressive ratio session.

### Fixed-Ratio Quinine + Alcohol Seeking

Mice remained on an FR4 operant schedule for quinine + alcohol seeking sessions. For these sessions, mice received liquid reinforcers of 9% ethanol containing 0-1000 µM quinine. Days 1-2 the reinforcer was 9% ethanol alone, Days 3-4 the reinforcer was 9% ethanol + 250 µM quinine, Days 5-6 the reinforcer was 9% ethanol + 500 µM quinine, and Days 7-8 the reinforcer was 9% ethanol + 1000 µM quinine.

*Chemogenetic inhibition during fixed-ratio quinine + alcohol seeking:* For 250 µM and 1000 µM quinine sessions, all mice were treated with both saline and CNO on consecutive days (3 mg/kg i.p., 30 minutes before the operant session) in a counterbalanced within-subject design.

### Tissue processing and viral expression validation

To prepare tissue for histology, mice were anesthetized with Avertin (1 ml, i.p.) and transcardially perfused with chilled 0.01 M phosphate-buffered saline (PBS) followed by 4% paraformaldehyde (PFA) in PBS. Brains were extracted and post-fixed in 4% PFA for 24h and then stored in PBS at 4°C for long-term storage. 45 µm coronal sections were collected using a Leica VT1000S vibratome (Leica Microsystems) and stored in 0.02% Sodium Azide (Sigma Aldrich) in PBS until histology was performed. BNST slices were mounted on slides and cover-slipped with Vectashield Hardset (Vector Biolabs) containing the nuclear counterstain DAPI. Viral infections were visualized using an epi-fluorescent microscope (Leica Microsystems) or a LSM 800 Confocal Microscope (Carl Zeiss, Inc.). Mice with viral infections outside of the BNST were excluded from the study.

### Chemogenetic inhibition + Immunohistochemistry (IHC)

To perform IHC following chemogenetic inhibition, mCherry and hM4di sucrose self-administration mice were injected i.p. with 3 mg/kg CNO and placed alone in a clean cage for two hours until tissue processing (see above section). To discourage behaviors that might create noise in our cFos measurements, we removed nesting material and plastic huts from these cages for the two-hour CNO period. For cFos IHC, tissue was washed for 3×10 minutes in PBS, permeabilized for 30 minutes in 0.5% Triton-X-100 in PBS, and immersed in blocking solution for one hour (0.1% Triton-X-100 + 10% Normal Donkey Serum in PBS). Next, the tissue was incubated overnight at 4°C in primary antibody diluted in blocking solution [rabbit anti-cFos 1:1000 (Synaptic Systems, Cat#20079)]. The next day, tissue was washed 4×10 minutes in PBS before being incubated for two hours in secondary antibody diluted in PBS (at 1:200) [Donkey anti-rabbit Cy2 (Jackson Immunoresearch, Cat#711-225-152)]. The tissue was then washed 3×10 minutes in PBS, mounted on slides, and allowed to dry overnight before cover slipping with Vecta-Shield Hardset Mounting Medium with DAPI (Vector Laboratories).

### Confocal Microscopy

All fluorescent IHC images were acquired with a Zeiss 800 upright confocal microscope using Zen Blue software (Carl Zeiss). Images were acquired using a 20x objective. Images were processed and quantified in FIJI [13] and MATLAB (Mathworks, Inc.).

### Statistics

Single-variable comparisons between two groups were made using paired or unpaired two-tailed t-tests where appropriate. If datasets had unequal variances, a non-parametric Mann-Whitney U-test was used. Grouped comparisons were made using two-way ANOVA or two-way mixed-model ANOVA (depending on the number of independent and within-subject variables), accounting for repeated measures when appropriate. Following significant interactions or main effects, appropriate post-hoc pairwise t-tests were performed using Tukey or Bonferroni post-hoc tests. All data are expressed as mean ± standard error of the mean (SEM), with statistical significance defined as p<0.05. All data were analyzed with GraphPad Prism 9 (GraphPad Software).

## Results

### Genetic knockdown of 5HT_2c_ receptors in the BNST does not strongly influence sweetened alcohol responding or motivation for sweetened alcohol in either sex, but may modestly influence quinine-adulterated alcohol seeking in females

Four weeks prior to the start of self-administration training, adult male and female 5HT2c*^lox/lox^* mice were stereotaxically injected with either AAV-Cre-GFP or AAV-GFP in the BNST and allowed to recover in their home cages. We have employed this transgenic mouse line in previous studies and have verified using *in-situ* hybridization that our AAV-Cre-GFP viral approach reduces 5HT_2c_ mRNA expression by ∼45% in the BNST [9]. All mice were group housed for the entirety of all experiments to prevent the influence of social isolation on alcohol consumption [14]. We restricted experimental mice to 85-90% of their original body weight and included sucrose in our alcohol solution (9% EtOH + 2% sucrose) (Supplemental Figure 1). We chose these conditions because (1) we believe sweetened alcohol more accurately models the conditions in which alcohol is voluntarily consumed in humans, especially in individuals with little alcohol drinking experience, and (2) it promotes high levels of voluntary operant alcohol intake.

Both males and females in control and 5HT_2c_ knockdown groups successfully acquired operant alcohol conditioning (Extended Data Figure 1-1). Females displayed higher operant responding rates and alcohol consumption in our model compared to males, which is consistent with previous reports (Figure 1B-G) [15, 16]. We observed no effect of BNST 5HT_2c_ deletion on FR4 responding for alcohol in either sex (effect of virus, females: p=0.2668, males: p=0.6666). This suggests that increased binge-like consumption of alcohol in females following BNST 5HT_2c_ deletion, as has been previously reported, does not occur when using an operant approach.

**Figure 1:**
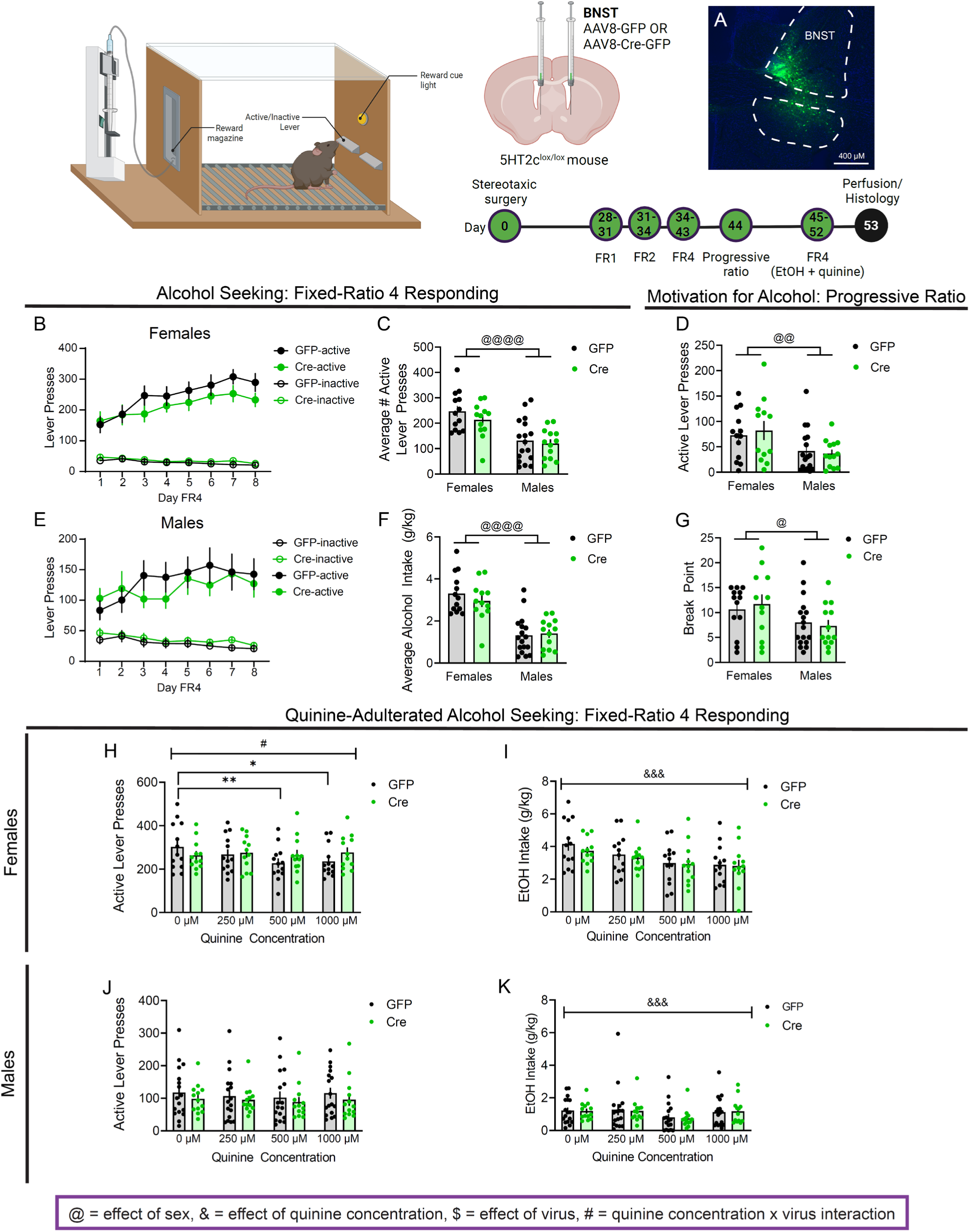
Genetic knockdown of the serotonin 5HT_2c_ receptor in the BNST promotes punished alcohol seeking in females. **A**, Representative viral infection in BNST. **B**, Active and inactive lever presses in FR4, females (Active: n=13 GFP, n=12 Cre; Two-way repeated measures ANOVA). **C**, Average active lever presses in FR4, both sexes (n=13 GFP females, n=12 Cre females, n=17 GFP males, n=13 Cre males; Two-way ANOVA). **D**, Active lever presses in PR, both sexes (n=13 GFP females, n=12 Cre females, n=17 GFP males, n=13 Cre males; Two-way ANOVA). **E**, Active and inactive lever presses in FR4, males (Active: n=17 GFP, n=13 Cre; Two-way ANOVA). **F**, Average alcohol intake in FR4, both sexes (n=13 GFP females, n=12 Cre females, n=17 GFP males, n=13 Cre males). **G**, Break point in PR, both sexes (n=13 GFP females, n=12 Cre females, n=17 GFP males, n=13 Cre males). **H**, Active lever presses for quinine-adulterated alcohol seeking, females (n=13 GFP, n=12 Cre; Two-way repeated measures ANOVA). **I**, Alcohol intake for quinine-adulterated alcohol seeking, females (n=13 GFP, n=12 Cre; Two-way repeated measures ANOVA). **J**, Active lever presses for quinine-adulterated alcohol seeking, males (n=17 GFP, n=13 Cre; Two-way repeated measures ANOVA). **K**, Alcohol intake for quinine-adulterated alcohol seeking, males (n=17 GFP, n=13 Cre; Two-way repeated measures ANOVA). # denotes quinine concentration x virus effect, & denotes effect of quinine concentration, @ denotes effect of sex. Figure made with Biorender.com. All data presented as mean ±SEM.

Next, to see if BNST 5HT_2c_ receptors regulate the motivation for alcohol, we tested the mice in a progressive ratio test session. In this test, the number of lever presses required to earn a single alcohol reward increased arithmetically over the session (see methods for details). This maximum number of alcohol rewards each animal was willing to work for was recorded as the “break point.” There were no differences between GFP and Cre mice in the progressive ratio test for either sex (Figure 1D, G) (effect of virus, active lever presses: p=0.8466, break point: p=0.9007). These data suggest that 5HT_2c_ in the BNST does not influence motivation for alcohol in either sex.

Following progressive ratio testing, we **removed** the sucrose from the alcohol solution used in operant conditioning and added in increasing amounts of the bitter tastant quinine (0 µM, 250 µM, 500 µM, 1000 µM). As quinine concentration increased, active lever presses and alcohol consumption were expected to plummet due to the unpleasant taste (aversive consequence). While females are generally less sensitive to the effects of quinine on operant responding for alcohol, this procedure has been widely implemented across the alcohol research field to investigate the negative consequences of alcohol consumption in both sexes (often referred to as compulsive or punished alcohol seeking) [15, 17–19]. Another benefit of this approach is that we were able to assess operant responding for alcohol in the **absence** of either sucrose or quinine for two days, thus helping inform potential confounds (0 µM condition). Once quinine was added to the alcohol solution, we did not include any sucrose in it. In males, BNST 5HT_2c_ knockdown did not affect active lever pressing for quinine-adulterated alcohol or the consumption of quinine-adulterated alcohol rewards (Figure 2J, K). We will note that neither GFP nor Cre males in this experiment reduced their operant responding as quinine concentration increased; however, consumption of quinine alcohol decreased with escalating quinine concentrations in both groups (effect of quinine concentration p<0.0001). The reasons for this disparity are not clear, but could speak to the development of habitual lever pressing after many weeks of training. GFP females reduced active lever presses and alcohol consumption with increasing quinine concentrations (Figure H, I) (GFP 0 µM vs. 500 µM: p=0.0039, GFP 0 µM vs. 1000 µM p=0.0123). However, we observed an interaction between quinine concentration and virus treatment such that Cre females did not reduce their active lever pressing for quinine alcohol as the quinine concentration increased. This effect was subtle—no post-hoc group differences at any quinine concentration were observed (quinine concentration x virus interaction p=0.0367). Together, these results suggest that 5HT_2c_ in the BNST does not robustly influence general alcohol self-administration behavior in males or females, but that it may modestly mediate aversive responses to alcohol in females.

**Figure 2:**
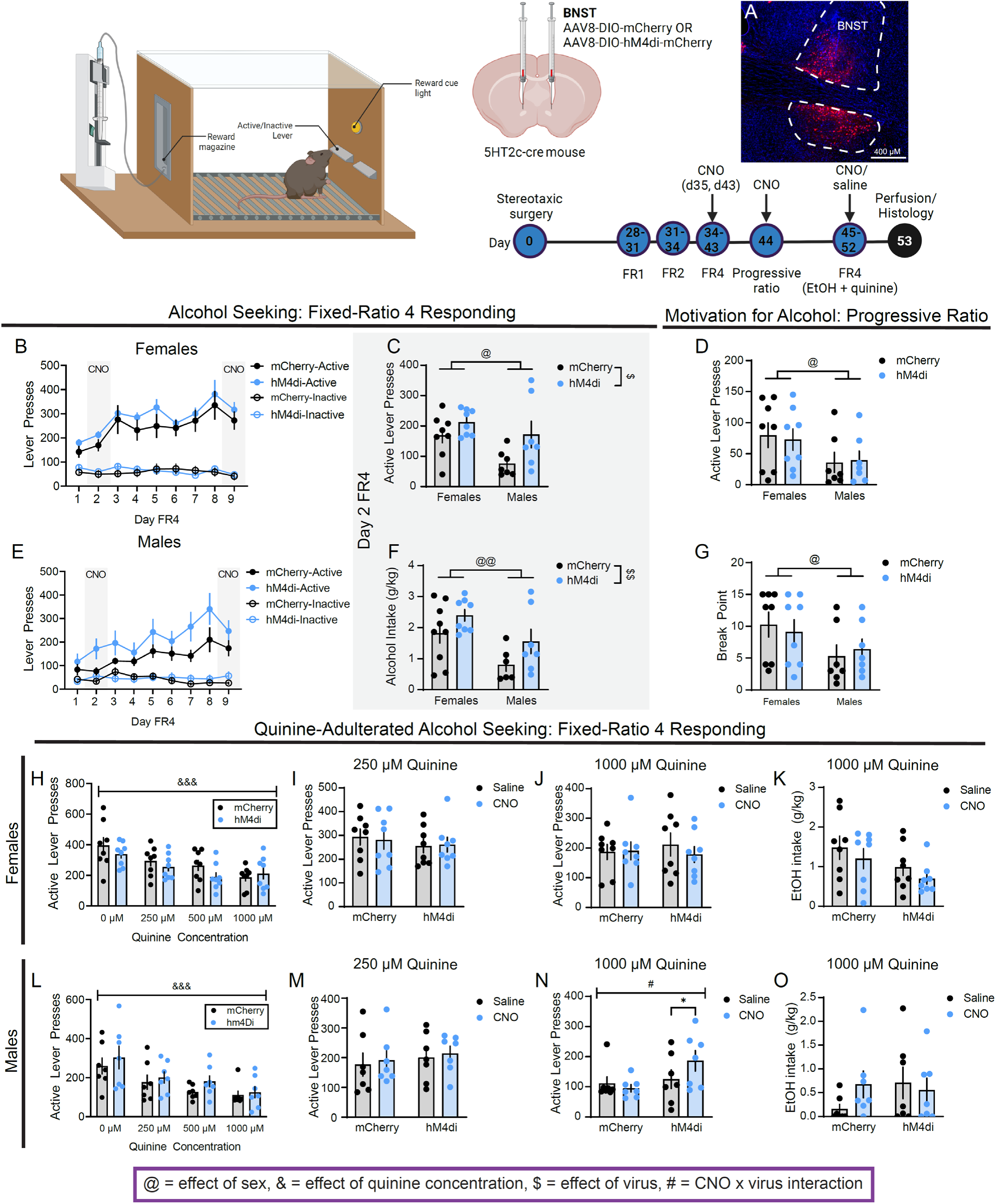
Chemogenetic inhibition of BNST^5HT2c^ promotes punished alcohol seeking in males. **A**, Representative image of viral infection in BNST. **B**, Lever presses in FR4, females (n=8 mCherry, n=8 hM4di; Two-way repeated measures ANOVA (d2-d8)). **C**, Active lever presses day 2 FR4 during chemogenetic inhibition, both sexes (n=8 mCherry females, n=8 hM4di females, n=7 mCherry males, n=7 hM4di males; Two- way ANOVA). **D**, Active lever presses in PR, both sexes (n=8 mCherry females, n=8 hM4di females, n=7 mCherry males, n=7 hM4di males). **E**, Lever presses in FR4, males (n=7 mCherry, n=7 hM4di; Two-way repeated measures ANOVA (d2-d8). **F**, Alcohol intake on d2 FR4 during chemogenetic inhibition, both sexes (n=8 mCherry females, n=8 hM4di females, n=7 mCherry males, n=7 hM4di males; Two-way ANOVA). **G**, Break point in PR, both sexes (n=8 mCherry females, n=8 hM4di females, n=7 mCherry males, n=7 hM4di males; Two- way ANOVA). **H**, Active lever presses for quinine-adulterated alcohol seeking without chemogenetic manipulation, females (n=8 mCherry, n=8 hM4di; Two-way repeated measures ANOVA). **I**, Active lever presses for alcohol + 250 uM quinine during chemogenetic inhibition, females (n=8 mCherry, n=8 hM4di; Two-way repeated measures ANOVA). **J**, Active lever presses for alcohol + 1000 uM quinine during chemogenetic inhibition, females (n=8 mCherry, n=8 hM4di; Two-way repeated measures ANOVA). **K**, Alcohol + 1000 uM quinine intake during chemogenetic inhibition, females (n=8 mCherry, n=8 hM4di; Two-way repeated measures ANOVA). **L**, Active lever presses for quinine-adulterated alcohol seeking without chemogenetic inhibition, males (n=7 mCherry, n=7 hM4di; Two-way repeated measures ANOVA). **M**, Active lever presses for alcohol + 250 uM quinine during chemogenetic inhibition, males (n=7 mCherry, n=7 hM4di; Two-way repeated measures ANOVA). **N**, Active lever presses for alcohol + 1000 uM quinine during chemogenetic inhibition, males (n=7 mCherry, n=7 hM4di; Two-way repeated measures ANOVA). **O**, Alcohol + 1000 uM quinine intake during chemogenetic inhibition, males (n=7 mCherry, n=7 hM4di; Two-way repeated measures ANOVA). $ denotes effect of virus, & denotes effect of quinine concentration, @ denotes effect of sex, # denotes CNO x virus interaction effect, * denotes post-hoc effects. Figure made with Biorender.com. All data presented as mean ±SEM.

### Genetic knockdown of 5HT_2c_ in the BNST reduces body weight selectively in males

We monitored mouse body weights daily during self-administration testing periods to ensure that experimental groups were equally food restricted compared to their relevant controls (85-90% of body weight at start of self-administration training). We discovered that 5HT_2c_ knockdown mice of both sexes weighed on average less than their control counterparts (females: effect of virus p=0.062; males: effect of virus p=0.0021) (Figure S1-1). While the difference in body weight between Cre and GFP females was observed on the day of surgery (p=0.0496), the difference in body weight between Cre and GFP males was not (p=0.8773). Therefore, these results suggest that the 5HT_2c_ signaling in the BNST is important for maintaining body weight in males, but not females.

### Chemogenetic inhibition of BNST^5HT2c^ increases operant responding for alcohol and alcohol intake in early (d2), but not late (d9), self-administration sessions

Adult male and female 5HT_2c_-cre mice were stereotaxically injected in the BNST with AAV-DIO-mCherry or AAV-DIO-hM4di-mCherry. At least four weeks later, mice were trained in the same alcohol self-administration paradigm employed above, again using a sweetened alcohol solution (9% EtOH + 2% sucrose). For chemogenetic manipulations during FR4 (days 2 and 9) and progressive ratio sessions (day 10), both mCherry and hM4di mice were treated with 3 mg/kg CNO intraperitoneally 30 minutes prior to testing. Importantly, previous studies have demonstrated that CNO itself does not impact alcohol intake in a home cage binge drinking model [9]. We found that chemogenetic inhibition of BNST^5HT2c^ on day 2, but not day 9, of FR4 self-administration increased active lever pressing (Day 2: effect of sex p=0.0184, effect of virus p=0.0148) and alcohol consumption (Day 2: effect of sex p=0.0047, effect of virus p=0.0369) in both sexes (Figure 2C, F, Extended Data Figure 1-1). While chemogenetic inhibition of BNST^5HT2c^ in males on day 2 appeared to increase operant alcohol self-administration on later days (Figure 2E, d2-d8), this effect was not statistically significant (effect of virus d2-d8 p=0.13). There was no effect of chemogenetic inhibition on progressive ratio responding in either sex (Figure 2D, G). These results indicate that BNST^5HT2c^ serve to constrain early FR4 operant responding for alcohol, and that this does not occur via increased motivation for alcohol.

### Chemogenetic inhibition of BNST^5HT2c^ promotes operant responding for quinine-adulterated alcohol in males

We next asked whether chemogenetic inhibition of BNST^5HT2c^ would influence operant responding for quinine-adulterated alcohol. For this set of experiments, we employed a different design than we did in our previous chemogenetics experiments. Rather than using a between-subjects design, here we used a within-subjects design. The rationale for this was that as mice are learning to operantly respond, as they were during the sweetened alcohol (no quinine) FR4 sessions, the number of responses reliably increases from one day to the next. This makes it difficult to accurately compare responding on successive days, which would be required for a within-subject design. This is not an issue during alcohol + quinine testing, as mice have stably learned the task and we observe no differences in lever pressing from one day to the next when considering a single quinine concentration. Weadministered either saline or 3 mg/kg CNO i.p. 30 minutes before the 250 µM and 1000 µM quinine sessions (counterbalanced for order of treatments). No sucrose was in the alcohol for the quinine adulterated alcohol self-administration experiments. When considering only the days where mice received no injections or only saline injections, both males and females reduced their operant responding for quinine alcohol as quinine concentrations increased (Figure 2H, L; males: effect of quinine concentration p<0.0001, females: effect of quinine concentration p<0.0001).Without CNO there were no differences observed between mCherry and hM4di groups of either sex. Chemogenetic inhibition of BNST^5HT2c^ did not affect operant responding for alcohol + 250 µM quinine in either sex (Figure 2I, M). However, this manipulation did increase responding for alcohol + 1000 µM quinine selectively in males (CNO x virus interaction p=0.0194, Bonferroni post-hoc hM4di CNO vs. saline p=0.021) (Figure 2N). Despite this, hM4di males did not consume more alcohol + 1000 µM quinine in the CNO condition compared to the saline condition (Figure 2O). These results suggest that BNST^5HT2c^ play a role in constraining operant responding for alcohol, but not necessarily alcohol consumption, in experienced male drinkers when it is associated with highly negative consequences.

### Chemogenetic inhibition of BNST^5HT2c^ does not influence operant responding for sucrose in either sex

Our alcohol self-administration experiments suggest that BNST^5HT2c^ function constrains early operant responding for alcohol. To address the confound of adding sucrose to our alcohol solution, we performed chemogenetic inhibition of BNST^5HT2c^ in a series of sucrose-only self-administration experiments. Adult male and female 5HT_2c_-cre mice were stereotaxically injected with AAV-DIO-mCherry or AAV-DIO-hM4di-mCherry in the BNST and allowed to recover in their home cages. At least four weeks later, mice began sucrose self-administration training. Similar to our alcohol self-administration experiments, we performed chemogenetic inhibition of BNST^5HT2c^ on days 2 and 9 of FR4 sessions and during progressive ratio sessions. We found no effect of chemogenetic inhibition of BNST^5HT2c^ on sucrose seeking on either day 2 or day 9 of FR4 (Figure 3B, C, E, F). We also found no effects of chemogenetic manipulation of BNST^5HT2c^ on motivation for sucrose in the progressive ratio test (effect of virus p=0.5316) (Figure 3D, G). Together, these results suggest that the role of BNST^5HT2c^ in constraining early operant responding for alcohol does not generalize to natural rewards like sucrose.

**Figure 3:**
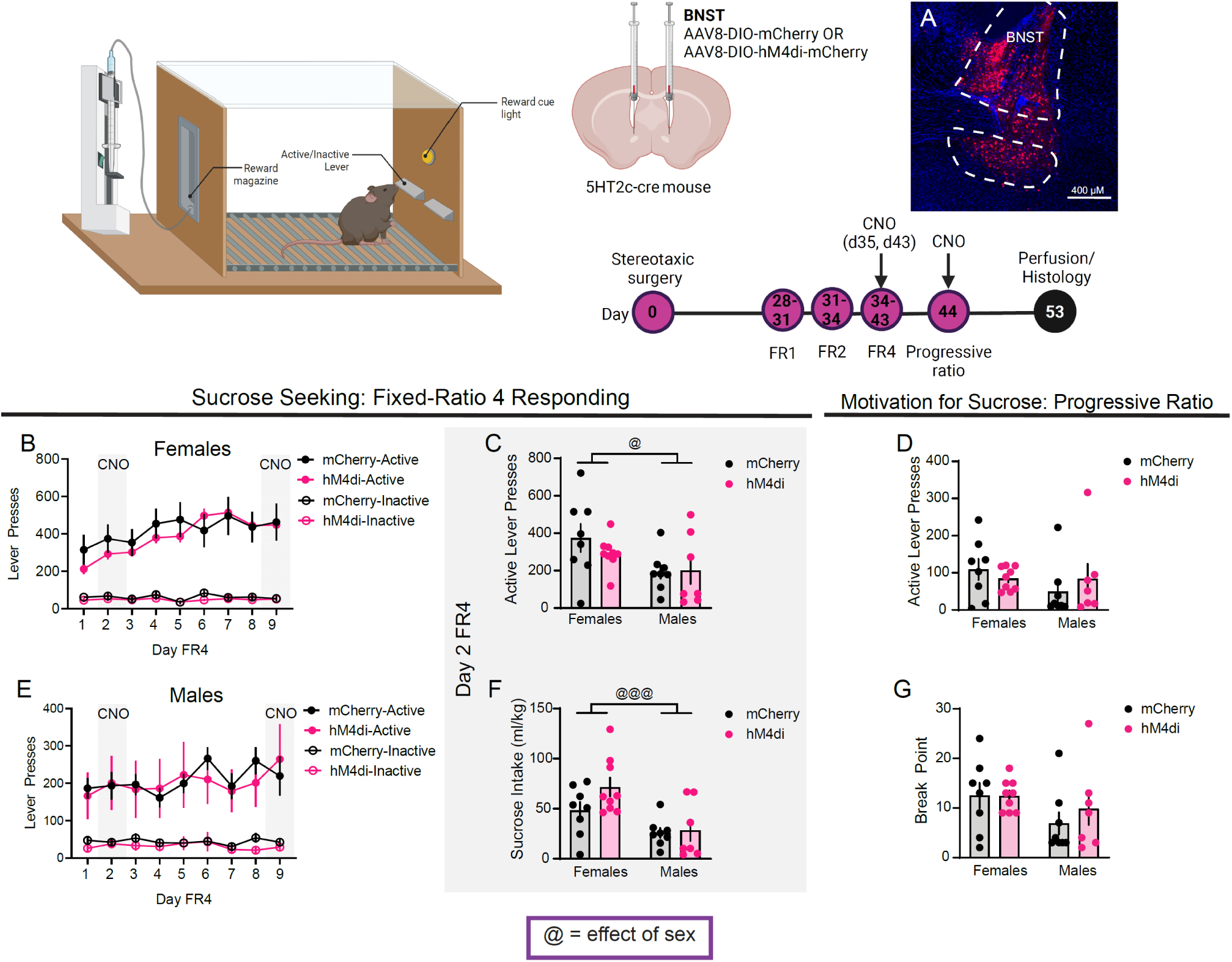
Chemogenetic inhibition of BNST^5HT2c^ does not influence sucrose seeking. **A**, Representative viral infection in the BNST. **B**, Lever presses in FR4 without chemogenetic inhibition, females (n=8 mCherry, n=9 hM4di). **C**, Active lever presses on d2 FR4 during chemogenetic inhibition, both sexes, all mice received 3 mg/kg CNO (n=8 mCherry females, n=9 hM4di females, n=8 mCherry males, n=7 hM4di males; Two-way ANOVA). **D**, Active lever presses in PR, both sexes during chemogenetic inhibition, both sexes, all mice received 3 mg/kg CNO (n=8 mCherry females, n=9 hM4di females, n=8 mCherry males, n=7 hM4di males; Two-way ANOVA). **E, L**ever presses in FR4 without chemogenetic inhibition, males (n=8 mCherry, n=7 hM4di). **F**, Sucrose intake on d2 FR4 during chemogenetic inhibition, all mice received 3 mg/kg CNO, both sexes (n=8 mCherry females, n=9 hM4di females, n=8 mCherry males, n=7 hM4di males; Two-way ANOVA). **G**, Break point in PR during chemogenetic inhibition, all mice received 3 mg/kg CNO, both sexes (n=8 mCherry females, n=9 hM4di females, n=8 mCherry males, n=7 hM4di males; Two-way ANOVA). @ denotes effect of sex. Figure made with Biorender.com. All data represented as mean ± SEM.

### BNST^5HT2c^ project to the basal forebrain, midbrain, and hindbrain

BNST^5HT2c^ have been reported to project to a variety of downstream regions [20]. However, a full characterization of BNST^5HT2c^ projection target regions in both sexes is lacking. This is especially important because understanding the anatomical connectivity of BNST^5HT2c^ neurons may help us isolate the neural circuit mechanisms mediating their specific roles in behavior. Thus, we performed anterograde tracing of BNST^5HT2c^ using a viral approach. We injected anterograde AAV-DIO-synaptophysin-mCherry into the BNST of 5HT2c-cre adult males and females (Figure 4A, B). This virus encodes for the presynaptic protein synaptophysin fused to an mCherry fluorescent tag, which allows us to visualize pre-synaptic terminal punctae in regions downstream from BNST^5HT2c^. We identified numerous putative synaptic targets of these neurons, all of which were shared among males and females: Nucleus Accumbens, Lateral Septum, Lateral Hypothalamus, Lateral Habenula, Ventromedial Hypothalamus, Central Amygdala, Ventral Tegmental Area, Dorsal Raphe Nucleus, and the Ventrolateral Periaqueductal Gray (Figure 4D-L).

**Figure 4:**
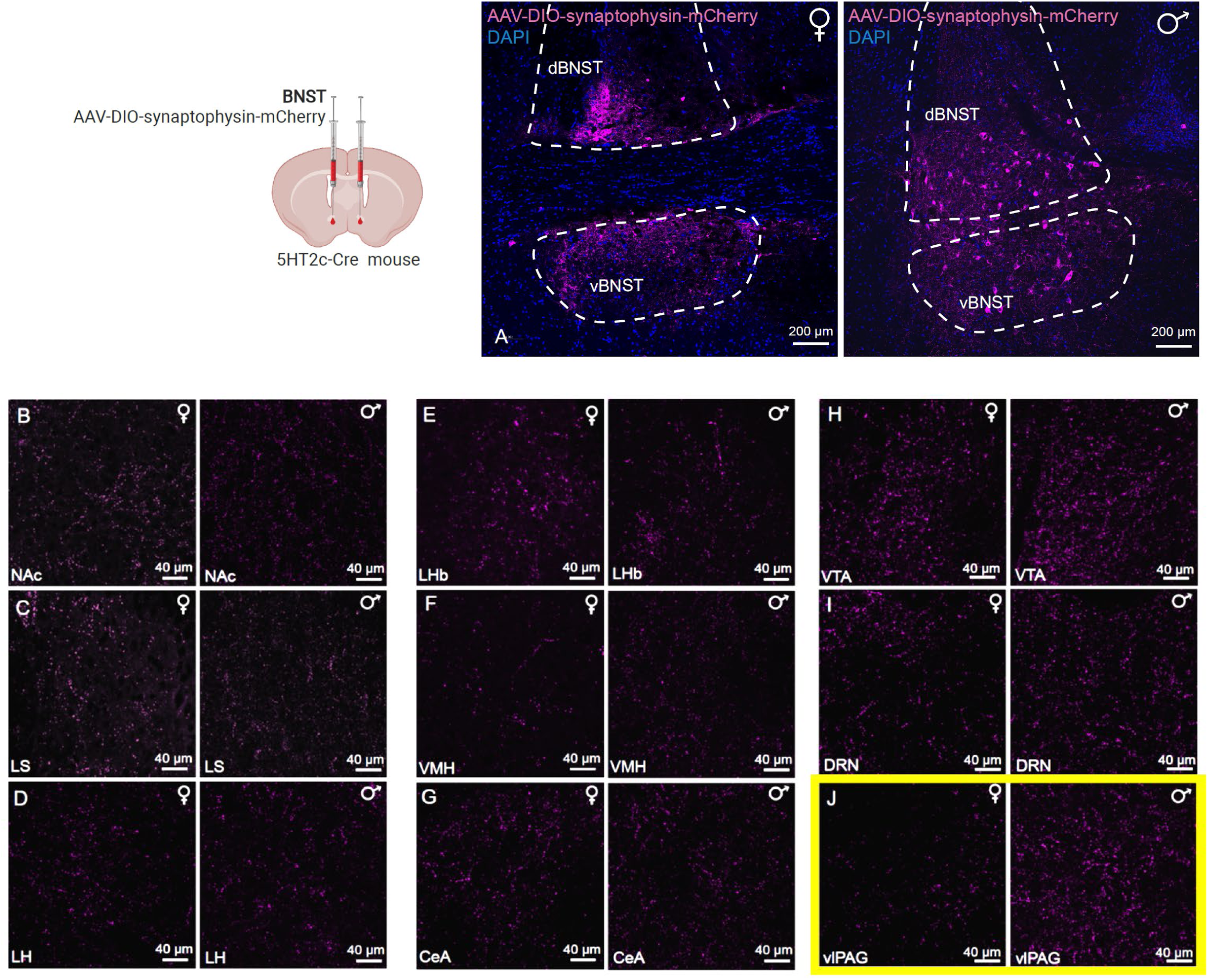
BNST5HT2c anterograde tracing in both sexes. A, Representative viral infections in BNST, females (left) and males (right). B, BNST^5HT2c^ post-synaptic terminals in Nucleus Accumbens (Nac). C, BNST^5HT2c^ post- synaptic terminals in Lateral Septum (LS). D, BNST^5HT2c^ post-synaptic terminals in Lateral Hypothalamus (LH). E, BNST^5HT2c^ post-synaptic terminals in Lateral Habenula (LHb). F, BNST^5HT2c^ post-synaptic terminals in Ventromedial Hypothalamus (VMH). G, BNST^5HT2c^ post-synaptic terminals in Central Amygdala (CeA). H, BNST^5HT2c^ post-synaptic terminals in Ventral Tegmental Area (VTA). I, BNST^5HT2c^ post-synaptic terminals in Dorsal Raphe Nucleus (DRN). J, BNST^5HT2c^ post-synaptic terminals in ventro-lateral Periacqueductal Gray (vlPAG); males show qualitatively more dense terminal expression than females.

### Chemogenetic inhibition of BNST^5HT2c^ reduces BNST and vlPAG activity in sex-dependent ways

The vlPAG has been previously implicated in aversive learning and consummatory behaviors, making it a potential candidate for mediating the behavioral effects of BNST^5HT2c^ inhibition on behavior. However, little is currently known about BNST projections to the vlPAG. To investigate how BNST^5HT2c^ might functionally influence vlPAG activity, we performed chemogenetic inhibition of BNST^5HT2c^ and stained for cFos in the vlPAG. One week after sucrose self-administration, we administered 3 mg/kg CNO i.p. to both mCherry and hM4di groups of both sexes and sacrificed the mice two hours later (no behavioral challenge). In order to appropriately compare DREADD-mediated effects in males and females, we first verified that both sexes had comparable expression of hM4di in the BNST (Figure 5A). As expected, acute chemogenetic inhibition of BNST^5HT2c^ reduced the number and percentage of virally-labeled BNST cFos+ neurons in both sexes (Figure 5D, E) (number: sex x virus interaction p=0.0452, virus p<0.0001, sex p=0.0312; percentage: sex x virus interaction p=0.0501, virus: p<0.0001, sex p=0.0022). Interestingly, we also observed suppression of total BNST cFos, which trended as more robust in males (Figure 5F) (sex x virus interaction p=0.0851, virus p=0.0082, sex p=0.0452). This could suggest that BNST^5HT2c^ exert more of an influence on local circuitry in males compared to females. Notably, both BNST and vlPAG activity were higher in males compared to females (main effects of sex p<0.05, Figure 5D, E, F, G, J, K). Chemogenetic inhibition of BNST^5HT2c^ reduced the number of cFos+ neurons in the vlPAG in both sexes (effect of virus p<0.0001) (Figure 5G-I), indicating that BNST^5HT2c^ disinhibit the vlPAG. As the anterior and posterior vlPAG have been reported to differ with respect to anatomy, connectivity, and behavioral function [21, 22], we also performed cFos counts at three different anterior-posterior vlPAG coordinates. Interestingly, DREADD inhibition of BNST^5HT2c^ reduced cFos primarily in the anterior vlPAG in females and primarily in the posterior vlPAG in males (Figure 5H, I). To directly compare the degree of disinhibitory control of BNST^5HT2c^ over vlPAG in males and females, we normalized the number of cFos+ vlPAG neurons to the number of DREADD- labeled neurons in the BNST for each animal (Figure 5J, K). This analysis revealed a trend for a larger total ratio in males (Figure 5J, p=0.0788). When considering anterior, middle, and posterior vlPAG separately, males showed a significantly higher ratio of vlPAG cFos to BNST DREADD cells (effect of sex: p=0.0058, Figure 5K). Together, these data indicate that when controlling for levels of DREADD expression, male BNST^5HT2c^ exert greater disinhibitory control over vlPAG neurons than female BNST^5HT2c^. This could potentially represent a downstream mechanism by which BNST^5HT2c^ inhibition permits increased operant alcohol responding for quinine alcohol in males.

**Figure 5:**
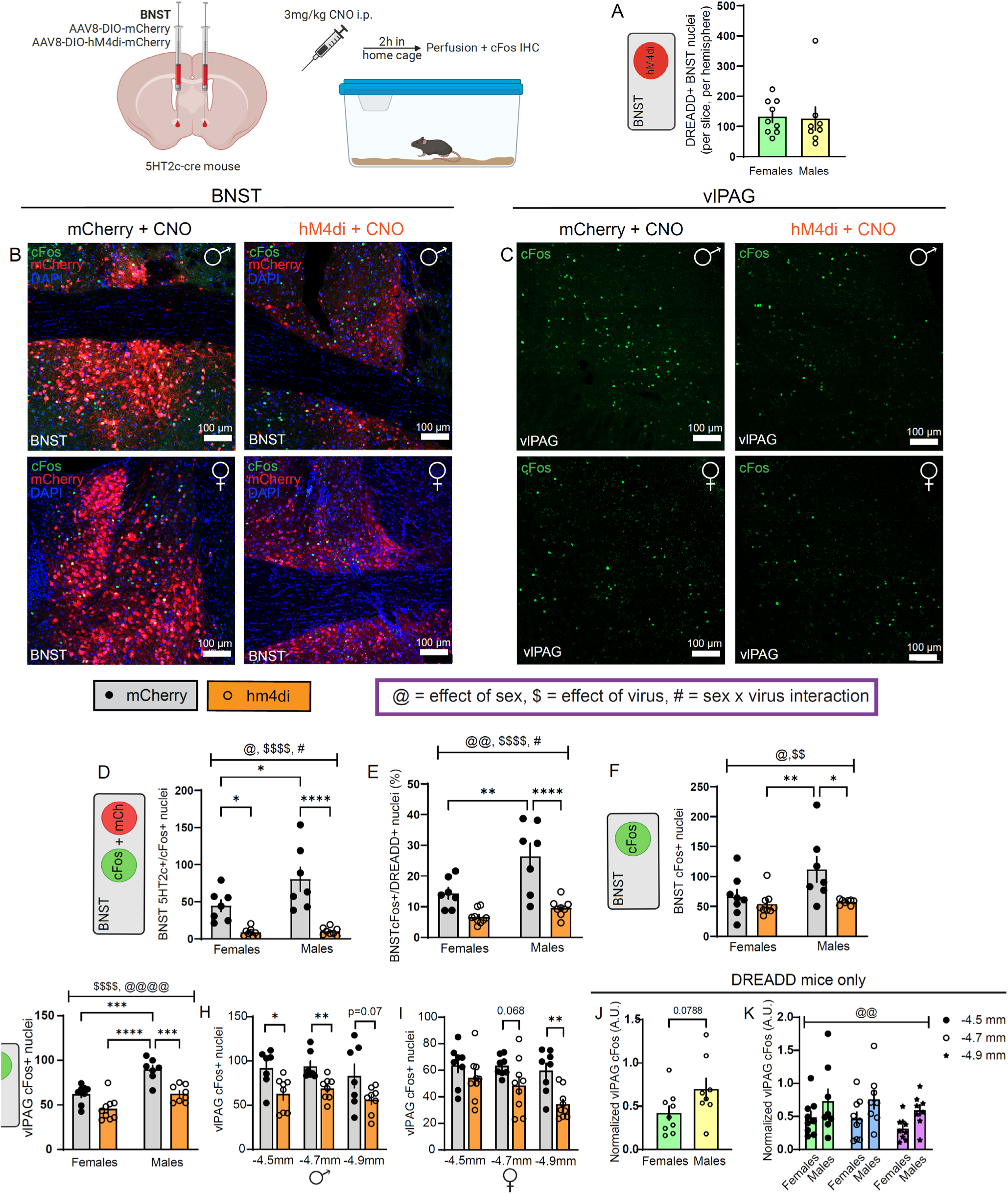
Chemogenetic inhibition of BNST^5HT2c^ reduces BNST and vlPAG activity in a sex-specific manner. **A**, Quantification of mCherry+ nuclei in DREADD (hM4di) males and females (n=9 females, n=8 males; Student’s two-tailed unpaired t-test). **B**, Immunohostochemistry for cFos (green), mCherry (red), and DAPI (blue) in the BNST following chemogenetic inhibition of BNST^5HT2c^. **C**, Immunohistochemistry for cFos (green) in the vlPAG following chemogenetic inhibition of BNST^5HT2c^. **D**, Quantification of BNST cFos+/5HT2c+ nuclei in both sexes (Two-way ANOVA and Bonferroni post-hoc; n=7 mCherry males, n=8 hM4di males, n=8 mCherry females, n=9 hM4di females). **E**, Quantification of BNST cFos as % of mcherry cells (Two-way ANOVA and Bonferroni posthoc, n=8 hM4di males, n=9 hM4di females). **F**, Quantification of total BNST cFos+ nuclei (Two-way ANOVA and Bonferroni post-hoc; n=7 mCherry male, n=8 hM4di male, n=8 mCherry female n=9 hM4di female). **G**, Quantification of vlPAG cFos+ neurons in males and females (n=7 mCherry male, n=8 hM4di male, n=8 mCherry female n=9 hM4di female). **H**, Quantification of vlPAG cFos+ nuclei in males (Student’s two-tailed unpaired t-tests; n=7 mCherry, n=8 hm4d). **I**, Quantification of vlPAG cFos+ nuclei in females (Student’s two-tailed unpaired t-tests; n=8 mCherry, n=9 hM4d). **J**, vlPAG cFos in males and females normalized to number of BNST mCherry cells in DREADD mice (# vlPAG cFos/# BNST mCherry) (Student’s two-tailed unpaired t-test; n=8 males, n=9 females **K**, vlPAG cFos in males and females normalized to number of BNST mCherry cells in DREADD mice (# vlPAG cFos/# BNST mCherry) (Two-way ANOVA, n=8 males, n=9 females). @ denotes effect of sex, $ denotes effect of virus, # denotes sex x virus interaction, * denotes post-hoc effect. Figure made with Biorender.com. All data represented as mean ± SEM.

## Discussion

Our results suggest that the 5HT_2c_ receptor in the BNST does not play a significant role in alcohol self-administration behaviors in males, and its role in alcohol self-administration in females is modest and may be specific to highly aversive outcomes linked to alcohol. However, the BNST^5HT2c^ neurons constrain the escalation of operant alcohol self-administration in both sexes. BNST^5HT2c^ project throughout the basal forebrain, midbrain, and hindbrain in both sexes, including the vlPAG. Functionally, BNST^5HT2c^ appear to disinhibit the vlPAG, which could possibly mediate the effects of BNST^5HT2c^ chemogenetic manipulation we observed on the alcohol self-administration behavior. Increased disinhibition of the vlPAG in males could possibly drive sex differences in the behavioral control of these neurons over responding for quinine-adulterated alcohol.

While the BNST has been reported to influence alcohol consumption in home cage models like Drinking in the Dark and Intermittent Access, few studies have investigated the role of this brain region in discrete aspects of alcohol consumption in a self-administration model. Recently, a study from our group found that knockdown of corticotropin releasing factor (CRF) peptide in the BNST reduced operant alcohol seeking and motivation for alcohol (Gianessi et al. 2023). Knockdown of the vesicular GABA transporter (vGAT) from BNST neurons expressing CRF (BNST^CRF^), however, produced the opposite effect, particularly in early FR4 sessions. Interestingly, CRF is released from a sub-population of BNST^5HT2c^ that are overwhelmingly GABAergic [8, 9]. Furthermore, 5HT_2c_ signaling itself influences CRF dynamics in the BNST: mice with knockout of 5HT_2c_ display reduced stress-induced activation of BNST^CRF^ neurons [11]. Functionally, knockdown of vGAT and chemogenetic inhibition of GABAergic neurons have common outcomes: reduced GABAergic output from these neurons. Given the overlap between BNST 5HT_2c_ and CRF systems, our present results are consistent with our previous self-administration study in BNST^CRF^ neurons, as we observed an increase in operant responding for alcohol in early, but not late, FR4 sessions with chemogenetic inhibition of BNST^5HT2c^.

An important question regarding our results is what an increase in early, but not late, operant alcohol self-administration signifies in terms of the behavioral processes impacted by BNST^5HT2c^ inhibition. We propose that the function of BNST^5HT2c^ in this context is to mediate the aversive effects of alcohol, which normally constrain early operant alcohol self-administration. These initially aversive qualities of alcohol primarily include unpleasant taste, motor impairment, and dysphoria [23]. However, these qualities are thought to dissipate with continued exposure to promote “aversion-resistant” drinking, particularly in female chronic drinkers [19]. In support of this idea that BNST^5HT2c^ are involved in the aversive effects of alcohol, we also observed an effect of BNST^5HT2c^ inhibition on responding for quinine alcohol in males at the highest concentration. While these mice did not consume more quinine alcohol than controls, they failed to reduce their responding when the alcohol was made highly aversive, suggesting some impairment in their behavioral responses to aversive outcomes related to alcohol. In addition, if BNST^5HT2c^ inhibition were increasing operant self-administration by increasing the rewarding effects of alcohol or its motivational salience, we would have likely observed increased responding on day 9 of fixed-ratio training and/or during the progressive ratio sessions.

The fact that we did not identify a robust role for the BNST 5HT_2c_ receptor in operant alcohol self-administration was contrary to our expectations based on previous work. However, we believe that the results of this study and a previous study [9], when considered together, may reveal a selective and sex-dependent role for the BNST 5HT_2c_ system in modulating the aversiveness of alcohol. We previously found using a drinking in the dark home cage binge model that knockdown of 5HT_2c_ in the BNST increased alcohol consumption selectively in females. In the current study, we observed only a subtle effect of this manipulation on quinine-adulterated alcohol responding, also selectively in females. There are multiple potential explanations for this discrepancy that do not preclude the conclusion that BNST 5HT_2c_ mediates, at least nominally, aversive responses to alcohol. First, if one considers the daily pattern of binge drinking in the home cage binge model in the prior study, the effect of BNST 5HT_2c_ knockdown on consumption was primarily driven by the first week of exposure. Indeed, by week three of drinking the control mice had escalated their consumption to match the knockdown mice, whose consumption stayed steady throughout the three weeks. This is consistent with overcoming the initially aversive qualities of alcohol, which become less important as one gains experience drinking. The effect of 5HT_2c_ knockdown on quinine-adulterated alcohol responding is consistent with this idea, but this effect was subtle. This subtlety could potentially be explained by the fact that the mice had such extensive experience self-administering alcohol over weeks that by the time quinine was introduced its aversiveness was partly overcome by habit or motivational drive. The addition of sucrose in our alcohol also may have contributed to an enhanced motivational drive for the alcohol that carried over to the quinine sessions (in which there was no sucrose in the alcohol). The potential evidence for this explanation is that our mice required a higher concentration of quinine to suppress their responding than what has been previously reported in other studies [15, 19, 24]. Similarly, the lack of effect of 5HT_2c_ knockdown on the early alcohol self-administration could be explained by the fact that the alcohol solution used here was likely less inherently aversive than the alcohol used in the previous home cage binge study. While the binge model exposed mice to 20% alcohol in water, our self-administration model used a 9% alcohol + 2% sucrose solution. Finally, and perhaps most importantly, mice were singly housed for the entirety of the home cage binge study while they remained group housed for the present operant self-administration studies. Social isolation is well known to have effects on mood, affect, motivation, and alcohol drinking behavior [14, 25, 26], and thus the stress of this housing condition could have influenced the manner in which BNST 5HT_2c_ knockdown affected alcohol consumption in Drinking in the Dark, particularly with regard to its initially aversive qualities. Overall, these results suggest that BNST 5HT_2c_ does not mediate alcohol consumption behavior in males, but in part mediates aversive responses to alcohol in females only when the alcohol is made sufficiently aversive.

We did not expect to observe effects of chemogenetic inhibition of BNST^5HT2c^ on alcohol self-administration behavior, as this manipulation previously had no influence on home cage binge drinking behavior in either sex. However, an important experimental condition to note from the previous study is that chemogenetic inhibition of BNST^5HT2c^ was performed on the last day of a three-week binge drinking period, which is likely after the time window where alcohol is aversive. Indeed, we found no effect in the present study of chemogenetic inhibition on day 9 of self-administration, but we observed an increase in self-administration on day 2. As mentioned above, it is possible that the length of our self-administration testing protocol combined with the addition of sucrose promoted high enough levels of operant responding such that higher concentrations of quinine were necessary to mediate strong response suppression in our model. This could potentially explain why we only observed an effect of chemogenetic inhibition at the highest concentration of quinine alcohol, rather than at both high and low concentrations in males. The fact that we did not observe this effect on quinine-adulterated alcohol responding in females could have been a consequence of a ceiling effect, as females performed more than double the number of active lever presses for 1000 uM quinine alcohol than males. Alternatively, this difference could reflect nuance in the level of control these neurons have over aversive behavioral responses to alcohol in males and females. For example, perhaps robust disinhibition of the vlPAG by BNST^5HT2c^ is required for aversive responses to alcohol in experienced drinkers such that inhibition of BNST^5HT2c^ in females is insufficient to alter this behavior. Therefore, the results of our present study and the previous study are in fact consistent with each other, and suggest a role for BNST^5HT2c^ in mediating initial aversive responses to alcohol in both sexes, and mediating aversive responses to alcohol’s consequences in experienced male drinkers. In addition, we want to note the importance of testing individual molecular and physiological manipulations in multiple models of voluntary alcohol consumption, as this allowed us to formulate a more refined picture of the role of the BNST 5HT_2c_ system in alcohol-related behaviors.

Our chemogenetic cFos mapping study suggests that BNST^5HT2c^ (which are GABAergic) disinhibit vlPAG by inhibiting the activity of vlPAG GABA neurons. This is consistent with a recent study in which BNST^GABA^ projections to vlPAG^GABA^ were functionally mapped using a combination of electrophysiology and optogenetics [22]. Notably, this study also found that BNST^GABA^-vlPAG^GABA^ projections enhance feeding behavior in male mice. However, given the specificity of our self-administration results and the fact that we did not observe increased self-administration of sucrose, it is unlikely that the findings in our chemogenetic inhibition experiments were the result of changes in feeding. On the other hand, our results do in fact suggest some role for the BNST 5HT_2c_ receptor in feeding behavior, as our male knockdown mice weighed less than their age-matched controls. Given that 5HT_2c_ is a Gq coupled receptor, this could confer a reduction in serotonin-mediated activation of BNST^GABA^ neurons, which could reduce feeding through reduced output to the vlPAG. But, in this case, if feeding were to influence sweetened alcohol self-administration, which is caloric, we would expect a reduction in feeding to correspond to a reduction in sweetened alcohol and/or sucrose self-administration (general reduction in consummatory/ingestive behavior). Future studies should determine directly whether and how the BNST 5HT_2c_ receptor, BNST^5HT2c^, and their projections to the vlPAG are involved in regulation of body-weight behavior, and whether that role is sex-dependent.

Food deprivation and the addition of sucrose to the alcohol solution are factors meant to encourage alcohol consumption and improve the face validity of our self-administration model. However, we recognize that these conditions introduce potentially confounding variables. To address the confound of sucrose in our sweetened alcohol self-administration experiments, we also performed chemogenetic inhibition of BNST^5HT2c^ in a sucrose-only self-administration model and observed no effects of our manipulations. However, sucrose is not aversive, even in inexperienced mice, and thus further experiments should be performed in future studies to test whether these neurons can increase punished sucrose self-administration (for example, punished with a shock). However, in support of the model we employed here, we would like to note that half of American adults consume sugar-sweetened beverages on a given day [27], and most commercially available flavored alcoholic beverages are sweetened with sugar [28]. With regard to the mild food restriction in our model, 1 in 5 Americans are actively dieting, which typically involves some level of food restriction [29]. In addition, intentional food restriction to enhance the pharmacological effects of alcohol consumption is common among binge drinkers and predictive of future alcohol-related issues [30]. Thus, while we recognize these caveats, we would also argue that the model we employed in this study is accurately reflective of common human behaviors and lifestyle factors.

In sum, we report that BNST 5HT_2c_ system (5HT_2c_ itself, BNST^5HT2c^ neurons) mediate some of the aversive effects of alcohol, especially relatively early on in an individual’s exposure. The role of 5HT_2c_ itself in this process appears modest and is specific to highly aversive alcohol-related outcomes in females. BNST^5HT2c^ appear to mediate initial aversion to alcohol in both sexes, while mediating aversion to alcohol-related negative outcomes in males with drinking experience. BNST^5HT2c^ neural projections could influence these behaviors through disinhibition of the vlPAG, which is more robustly modulated by BNST^5HT2c^ in males than in females. These findings may have implications for the development of treatments for AUD and other disorders of impulsivity.

## Supporting information

Statistical Table Flanigan et al 2023b

## Acknowledgements

Figures for this manuscript were created using BioRender.com. The authors would like to acknowledge Drs. Alison Roland and Brianna George for their assistance editing this manuscript.

## Author Contributions

M.E.F., C.G., and T.L.K. conceived of and designed experiments. M.E.F., C.G., M.C., W.D., and V.S. performed behavioral experiments. M.F., M.C. and V.S. performed histology. C.G. and M.E.F. performed intracranial surgeries. M.E.F. analyzed data and wrote manuscript with edits from all authors.

---

**Extended Data Figure 1-1:**
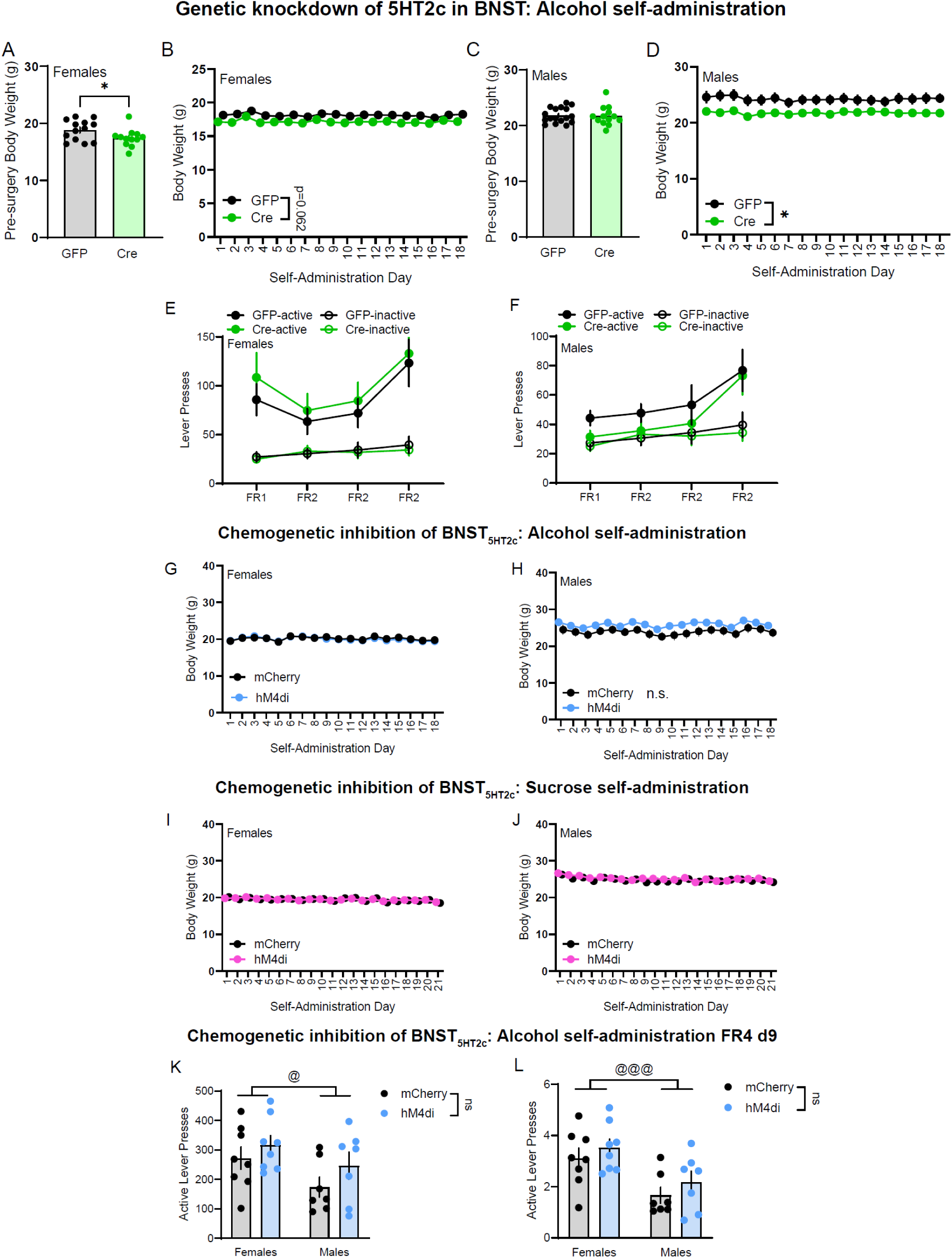
Body weights for all cohorts; Additional chemogenetic inhibition data. **A**, Pre-surgery body weights, females (n=13 GFP, n=12 Cre, Student’s two-tailed unpaired t-test, t(23)=2.072, p=0.0496. **B**, Genetic knockdown of 5HT2c in the BNST reduces body weight in females (n=13 GFP, n=12 Cre; Two-way repeated measures ANOVA; virus x day F(17,374)=0.3202, p=0.9959; day F(17.374)=4.292, p<0.0001; virus F(1,22)=3.926, p=0.062). **C**, Pre-surgery body weights, males (n=17 GFP, n=13 Cre, Student’s two-tailed unpaired t-test, t(28)=0.1558, p=0.8773). **D**, Genetic knockdown of 5HT2c in the BNST reduces body weight in males (n=17 GFP, n=13 Cre; Two-way repeated measures ANOVA; virus x day F(17,476)=1.518, p=0.0835; day F(17,476)=3.790, p<0.0001; virus F(1,28)=4.820, p=0.0366). **E**, Genetic knockdown of 5HT2c in the BNST does not influence the acquisition of operant responding for alcohol in females (n=13 GFP, n=12 Cre, Two-way repeated measures ANOVA, day x virus: F(3,72)0.1218, p=0.9470; day: F(3,72)=9.409, p=0.0002; virus: F(1,24)=0.3658, p=0.5510). **F**, Genetic knockdown of 5HT2c in the BNST does not influence the acquisition of operant responding for alcohol in males (n=17 GFP, n=13 Cre; Two-way repeated measures ANOVA; day x virus: F(3,84)=0.1331, p=0.9401; day: F(3,84)=7.349, p=0.0002; virus: F(1,28)=1.227, p=0.2775). **G**, Body weights did not differ between mCherry and hM4di groups for alcohol self-administration, females (n=8 mCherry, n=8 hM4di, Two-way repeated measures ANOVA; day x virus F(17,238)=2.104, p=0.0076; day F(17,238)=23.37, p<0.0001; virus F(1,14)=0.03912, p=0.8461). **H**, Body weights did not differ between mCherry and hM4di groups for alcohol self-administration, males (n=7 GFP, n=7 Cre; Two-way repeated measures ANOVA; day x virus F(17, 204)=1.577, p=0.0727; day F(17,204)=26.51, p<0.0001; virus (1,12)=2.244, p=0.1599). **I**, Body weights did not differ between mCherry and hM4di groups for sucrose self-administration, females (n=8 GFP, n=10 Cre, Two-way repeated measures ANOVA; virus x day F(20, 320)=1.882, p=0.0131; day F(20,320)=15.78, p<0.0001; virus F(1,16)=0.01885, p=0.8925). **J**, Body weights did not differ between mCherry and hM4di groups for sucrose self-administration, males (n=8 GFP, n=7 Cre, Two-way repeated measures ANOVA; virus x day F(20,250)=2.924, p<0.0001; day F(20,260)=20.40, p<0.0001; virus F(1,13)=0.1819, p=0.6768). **K**, Chemogenetic inhibition of BNST^5HT2c^ does not alter the maintenance of alcohol seeking behavior (n=8 mCherry females, n=8 hM4di females, n=7 mCherry males, n=7 hM4di males; Two-way ANOVA; sex x virus F(1,26)0.1438, p=0.7076; sex F(1,26)=5.025, p=0.0337; virus F(1,26)=2.453, p=0.1264). **L**, Chemogenetic inhibition of BNST^5HT2c^ does not alter alcohol intake on day 9 of self-administration (Two-way ANOVA, n=8 mCherry females, n=8 hM4di females, n=7 mCherry males, n=7 hM4di males; sex x virus F(1,26)=0.02165, p=0.884; sex F(1,26)=14.44 p=0.0008; virus F(1,26)=1.592 p=0.2182). @ denotes effect of sex, *p<0.05. Figure made with Biorender.com. All data represented as mean ± SEM.

## References

1. Schmidt, R.A., et al., The early impact of COVID-19 on the incidence, prevalence, and severity of alcohol use and other drugs: A systematic review. Drug Alcohol Depend, 2021. 228: p. 109065.

2. Kerr, W.C., et al., Longitudinal assessment of drinking changes during the pandemic: The 2021 COVID-19 follow-up study to the 2019 to 2020 National Alcohol Survey. Alcohol Clin Exp Res, 2022. 46(6): p. 1050–1061.

3. Bacqué-Cazenave, J., et al., Serotonin in Animal Cognition and Behavior. International Journal of Molecular Sciences, 2020. 21(5): p. 1649.

4. Hammack, S.E., et al., The response of neurons in the bed nucleus of the stria terminalis to serotonin: Implications for anxiety. Progress in Neuro-Psychopharmacology and Biological Psychiatry, 2009. 33(8): p. 1309–1320.

5. Guo, J.D., et al., Bi-directional modulation of bed nucleus of stria terminalis neurons by 5-HT: molecular expression and functional properties of excitatory 5-HT receptor subtypes. Neuroscience, 2009. 164(4): p. 1776–93.

6. Hazra, R., et al., Differential distribution of serotonin receptor subtypes in BNST(ALG) neurons: modulation by unpredictable shock stress. Neuroscience, 2012. 225: p. 9–21.

7. Marcinkiewcz, C.A., et al., Sex-Dependent Modulation of Anxiety and Fear by 5-HT1A Receptors in the Bed Nucleus of the Stria Terminalis. ACS Chemical Neuroscience, 2019. 10(7): p. 3154–3166.

8. Mazzone, C.M., et al., Acute engagement of G(q)-mediated signaling in the bed nucleus of the stria terminalis induces anxiety-like behavior. Mol Psychiatry, 2018. 23(1): p. 143–153.

9. Flanigan, M.E., et al., Subcortical serotonin 5HT2c receptor-containing neurons sex-specifically regulate binge-like alcohol consumption, social, and arousal behaviors in mice. Nature Communications, 2023. 14(1): p. 1800.

10. Marcinkiewcz, C.A., et al., Ethanol induced adaptations in 5-HT2c receptor signaling in the bed nucleus of the stria terminalis: implications for anxiety during ethanol withdrawal. Neuropharmacology, 2015. 89: p. 157–67.

11. Heisler, L.K., et al., Serotonin 5-HT2C receptors regulate anxiety-like behavior. Genes, Brain and Behavior, 2007. 6(5): p. 491–496.

12. Berglund, E.D., et al., Serotonin 2C receptors in pro-opiomelanocortin neurons regulate energy and glucose homeostasis. The Journal of Clinical Investigation, 2013. 123(12): p. 5061–5070.

13. Schindelin, J., et al., Fiji: an open-source platform for biological-image analysis. Nat Methods, 2012. 9(7): p. 676–82.

14. Lopez, M.F. and K. Laber, Impact of social isolation and enriched environment during adolescence on voluntary ethanol intake and anxiety in C57BL/6J mice. Physiol Behav, 2015. 148: p. 151–6.

15. Sneddon, E.A., et al., Increased responding for alcohol and resistance to aversion in female mice. Alcoholism: Clinical and Experimental Research, 2020. 44(7): p. 1400–1409.

16. Gianessi, C.A., et al., Disentangling the effects of Corticotrophin Releasing Factor and GABA release from the ventral bed nucleus of the stria terminalis on ethanol self-administration in mice. bioRxiv, 2023: p. 2023.03.02.530838.

17. Radke, A.K., et al., Recent perspectives on sex differences in compulsion-like and binge alcohol drinking. International Journal of Molecular Sciences, 2021. 22(7): p. 3788.

18. Radke, A.K., E.A. Sneddon, and S.C. Monroe, Studying sex differences in rodent models of addictive behavior. Current Protocols, 2021. 1(4): p. e119.

19. Sneddon, E.A., R.D. White, and A.K. Radke, Sex differences in binge-like and aversion-resistant alcohol drinking in C57 BL/6J mice. Alcoholism: clinical and experimental research, 2019. 43(2): p. 243–249.

20. Marcinkiewcz, C.A., et al., Serotonin engages an anxiety and fear-promoting circuit in the extended amygdala. Nature, 2016. 537(7618): p. 97–101.

21. Satpute, A.B., et al., Identification of discrete functional subregions of the human periaqueductal gray. Proceedings of the National Academy of Sciences, 2013. 110(42): p. 17101–17106.

22. Hao, S., et al., The Lateral Hypothalamic and BNST GABAergic Projections to the Anterior Ventrolateral Periaqueductal Gray Regulate Feeding. Cell Rep, 2019. 28(3): p. 616–624.e5.

23. Verendeev, A. and A.L. Riley, The role of the aversive effects of drugs in self-administration: assessing the balance of reward and aversion in drug-taking behavior. Behavioural pharmacology, 2013. 24(5 and 6): p. 363–374.

24. Siciliano, C.A., et al., A cortical-brainstem circuit predicts and governs compulsive alcohol drinking. Science, 2019. 366(6468): p. 1008–1012.

25. Lopez, M.F., T.L. Doremus-Fitzwater, and H.C. Becker, Chronic social isolation and chronic variable stress during early development induce later elevated ethanol intake in adult C57BL/6J mice. Alcohol, 2011. 45(4): p. 355–64.

26. Rivera-Irizarry, J.K., M.J. Skelly, and K.E. Pleil, Social Isolation Stress in Adolescence, but not Adulthood, Produces Hypersocial Behavior in Adult Male and Female C57BL/6J Mice. Front Behav Neurosci, 2020. 14: p. 129.

27. Rosinger, A., et al., Sugar-sweetened beverage consumption among US youth, 2011-2014. 2017.

28. Wakabayashi, K.T., et al., The Sugars in Alcohol Cocktails Matter. ACS Chem Neurosci, 2021. 12(18): p. 3284–3287.

29. Stierman, B., et al., Special diets among adults: United States, 2015-2018. 2020: US Department of Health and Human Services, Centers for Disease Control and ….

30. Simons, R.M., et al., Drunkorexia: Normative behavior or gateway to alcohol and eating pathology? Addictive Behaviors, 2021. 112: p. 106577.

